# The effect of particle size on wheat bran fermentation by human gut microbiota

**DOI:** 10.1101/2025.05.15.654185

**Authors:** Ashwana D. Fricker, Dane G. Deemer, Yunus E. Tuncil, Riya Thakkar, Arianna R. Marcia, Connor W. Emsley, Stephen R. Lindemann

## Abstract

0.

Dietary fibers within whole grains reach the large intestine where they shape the microbial composition. However, the bioavailability of these dietary nutrients to the microbiota is likely limited due to entrapment within the grain particle and requires liberation by microbial enzymes. Here, we used batch fecal fermentation from mixed donors on a range of sizes of wheat particles generated by cyclone milling from a single source to identify bacterial taxa and genomic signatures that are responsive to differences in wheat bran fine structures. We present evidence that different taxa within the same genus colonize wheat bran particles of different sizes. Further, neutral sugar content varied across wheat bran particles despite originating from the same batch, suggesting different polysaccharide structures and nutritional niches. In line with the taxonomic and compositional differences, specific short chain fatty acids varied across particle sizes; in fine wheat bran particle fermentations propionate was high and butyrate low. To identify relevant genomic features implicated in bran colonization, we took a metagenomic approach. From this, we linked genes associated with polysaccharide fermentation to wheat bran particles independent of size, however, within one well-distributed taxon, *Lachnospiraceae*, genes related to motility were linked to large and medium wheat bran particles. Overall, these results suggest that differences in fine structures and resource availability, as generated through milling, can drive compositional changes in the gut microbiota in an organism-specific manner, mediated through its genomic capacity.

**IMPORTANCE:** Cereal brans comprise a large fraction of the dietary fiber consumption. Although it is well-known that dietary fibers influence the metabolic output and taxonomic composition of the gut microbiota, relatively little is known regarding whether the fine structures and resource availability of milled whole grains exert any influence on the microbial makeup. Our data suggest that the sugar content varies across bran milled from a single source to different sizes. These differences in composition may result in colonization differences by related, but unique, taxa, mediated by genes related to polysaccharide fermentation, thus leading to differences in metabolic output. Furthermore, our data suggest that genes related to motility might influence the capacity of microorganisms to colonize particles. Taken together, our data suggest that physical context can influence gut microbiota composition in turn impacting metabolic output.

**Graphical Abstract:** 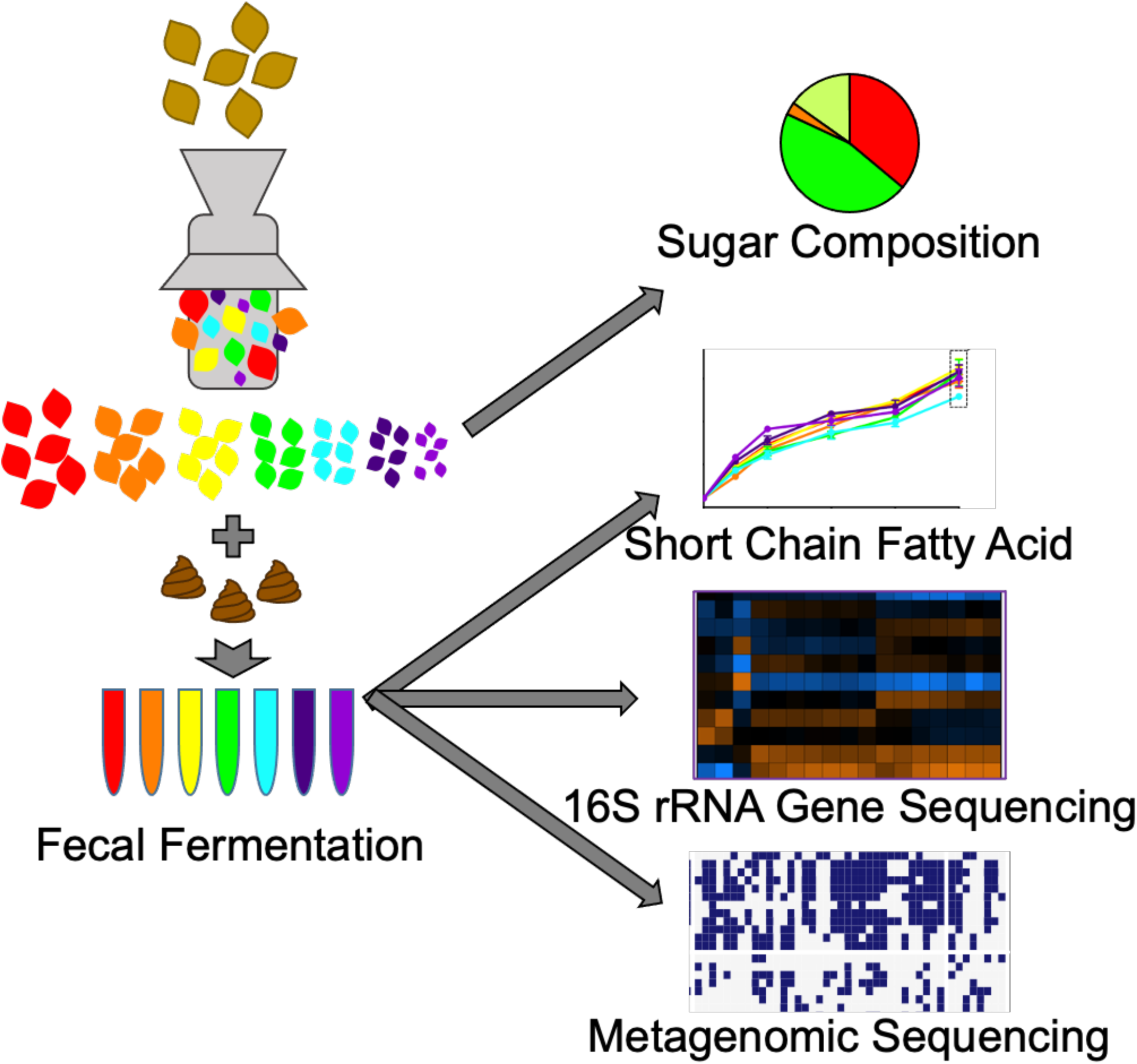

## 1. INTRODUCTION

The gut microbiome supports many aspects of human well-being, including food digestion (1), immune system support (2), and vitamin production (3). Diverse gut microbiomes are inversely correlated with diabetes, obesity, and inflammatory bowel disease, among others; however, a direct link between microbial community composition, diversity, and disease states remains elusive (4–10). One proposed mechanism for microbiome-induced health benefits is dietary fiber fermentation fibers into short-chain fatty acids (SCFAs), the terminal products of polysaccharide metabolism (11, 12). Each SCFA (acetate, propionate, and butyrate) provides distinct benefits depending on its extent of absorption and location of action (13). Although some functions are retained across taxonomically divergent communities (14), others are unique to specific taxa; consequently differences in community composition result in distinct determinants of fermentative functions which may degrade divergent fiber structures to distinct metabolites (15). Divergent fiber structures select for distinct microbial species (16–19), genotypes of which may vary across hosts via differences in genome content (20), mutations (21), or phage pressure (22) that translate to divergent growth patterns (23) and metabolic outputs (24). Such interrelationships illuminate the need to better characterize relationships among fine fiber structures, microbial genotypes, and metabolic outputs.

Approximately 50% of North American dietary fiber intake is from cereal brans, the rest of which is comprised of vegetables, legumes, fruits, and other sources (25). Wheat bran (WB) is a byproduct of the wheat milling and refining process (i.e. white flour production) and the dominant cereal bran in the U.S. diet. WB is rich in strongly fermented dietary fibers, (e.g., arabinoxylans and beta-glucans) and relatively low in poorly fermented fibers (e.g., cellulose and lignin) (26). Recent studies reveal that differences in dietary fiber chemical structure influence growth rates of individual species, given differences in linkage composition (16), arabinose : xylose ratio (27), degree of polymerization (28), and branch structure (19), and, for insoluble substrates, differences in particle size (17, 29).

Community metabolism of fiber structures is an emergent property driven by population-level efficiencies in degradation, internalization, and metabolism (generally considered at the level of species, although intraspecies variation can also be impactful) and their competitive and cooperative interactions (30). In complex structures, such as WB, access to individual dietary fibers is limited by their heterogeneous deposition in plant cell walls across tissue types (31, 32); degradation may require sequential activity of multiple microbial enzymes and/or result in resource patchiness and uneven liberation from particles (23). Patterns of differential spatial and temporal colonization arise in many complex systems including marine particles (33), lignocellulose (34), and in *in vitro* fermentation of maize (29) and wheat brans (17, 35). In these studies, individual operational taxonomic units (OTUs) classified in the same genus display different succession on differently sized particles (17, 29).

Here, we demonstrate that organismal gene content is associated with the ability to colonize variously sized WB particles. We hypothesized that genes involved in motility and attachment as well as polysaccharide fermentation would associate with WB colonization. We first took an amplicon sequencing approach to identify closely related species with unique growth patterns across bran sizes. We then characterized microbial metabolic networks related to dietary fiber fermentations and identified microbial functions that are tightly associated with particle physiochemical attributes. These findings highlight the importance of WB milling processes on the growth and metabolism of gut microorganisms.

## 2. METHODS

### 2.1 Wheat bran fractions collection and testing

WB was a gift of the Mennel Milling Company (Fostoria, OH). First, the largest bran fraction (i.e., > 1700 µm) was sieved out of mixed bran sample using a Portable Sieve Shaker Model RX-24 sieving machine with sieves of 180, 250, 300, 500, 850, 1000, and 1700 µm screen size (W. S. Tyler Combustion Engineering, Inc., Mentor, OH, U.S.). These were then milled using a cyclone mill and sieved again. Seven total WB size fractions were obtained: (1) 180–250 (2) 250–300 (3) 300–500 (4) 500–850 (5) 850–1000 (6) 1000–1700 and (7) >1700 μm as above. These seven size fractions were used for further experiments.

Each of these WB particle fractions were passaged through upper GI *in vitro* digestion as previously described (as Method A) (36). Briefly, WB fractions were treated with pepsin at pH 2.5 ± 0.1 to mimic conditions in the stomach for 30 min, pH was adjusted to 6.9 ± 0.1 and pancreatin and amyloglucosidase were added to mimic small intestinal passage for 6 h. This was followed by dialysis for 36 h and freeze drying prior to *in vitro* fermentation, as described.

Neutral monosaccharides from different WB fractions post-upper GI tract digestion were detected using gas chromatography (with helium as carrier gas) on a capillary column (SP2330, Supelco, Bellefonte, PA, U.S.) coupled with mass spectroscopy (GC–MS; 7890A gas chromatograph and 5975C inert mass selective detector, Agilent Technologies, Inc., Santa Clara, CA, U.S.) as described previously (36).

### 2.2 Fermentation and sample collection

Each WB fraction was weighed (44 ± 1 mg) into Balch tubes (Chemglass Life Sciences, Vineland, NJ, U.S.). Bottles containing phosphate-buffered gut mineral medium and tubes containing WB particles, fructooligosachharides (FOS; Sigma-Aldrich, St. Louis, MO, U.S. - positive control), or no carbon source (blank – negative control) were placed in the anaerobic chamber (BACTRONEX; Shel Lab, Cornelius, OR, U.S.) under an 90% N_2_, 5% CO_2_, and 5% H_2_ atmosphere overnight to remove oxygen. The phosphate-buffered gut mineral medium contained resazurin as an oxygen indicator and trace elements (8.0 mM NaCl, 6.3 mM KCl, 3.3 mM urea, 3.3 mM NH4Cl, 0.7 mM Na2SO4, 40 mM sodium phosphate buffer (pH 7.0), 1 mg resazurin, 0.25 g/L cysteine HCl, 333 μM CaCl2, 492 μM MgCl2, and 1X P1 metals and trace elements) (29).

On the day of inoculation, 4 ml of gut mineral medium was added to Balch tubes containing different WB fractions, the blank tubes, and the fermentation positive control (FOS). Fecal samples from three healthy donors who were consuming their routine diets and had not taken antibiotics for at least 3 months were pooled as previously described (17). To prevent the loss of bacterial viability, fecal samples were collected, maintained on ice, rapidly transferred to the anaerobic chamber, and used within 2 h of donation. Briefly, fecal samples were mixed with gut mineral media in the ratio 1:10 (w/v) and filtered through four layers of cheese-cloth. After filtration, fecal slurries from individual donors were pooled equally (37). Each Balch tube prepared as above then received 0.4 mL of the pooled fecal slurry mix, was sealed with butyl rubber stoppers, and crimped with aluminium seals (both from Chemglass Life Sciences, Vineland, NJ, U.S.), These were performed in triplicate and incubated at 37°C in a shaking incubator (Innova 42, New Brunswick Scientific, Edison, NJ, U.S.) at 150 rpm. Human stool collection and use protocols were reviewed and approved by Purdue University’s Institutional Review Board (IRB #1701018645).

At five time points (6, 12, 24, 36, and 48 h) post inoculation, we measured gas production, pH, and short-chain fatty acid (SCFA) concentrations. Gas production was measured as overpressure by passing a needle on a glass graduated syringe through the rubber stopper prior to unsealing the tubes. Supernatant pH was measured using a benchtop pH meter. At each time point, we collected a 0.4 mL aliquot from each tube for SCFA measurements to which an internal standard (157.5 μl of 4-methyl valeric acid, 1.47 ml of 85% phosphoric acid, 39 mg of copper sulfate pentahydrate in a total volume of 25 ml) was immediately added. Samples were then stored at −80°C.

SCFA measurements were made as previously described (38). Briefly, 4 μl of the supernatants were analyzed on a fused-silica capillary column (NukonTM, SUPELCO No: 40369-03A, Bellefonte, PA, U.S.) using a gas chromatograph (GC-FID 7890A, Agilent Technologies, Inc.) (38). Analytical grade acetate, propionate, and butyrate (Fisher Scientific, Hampton, NH, U.S.) were used as external standards.

At 12, 24, and 48 h post-inoculation, particles in tubes allowed to settle for ∼30 seconds. Thereafter, supernatants were carefully removed to separate the supernatant from fermented WB particles to minimize mixing of particle- and supernatant-associated microbes and stored at - 80°C.

### 2.3 DNA extraction and 16S (v4-v5) amplicon sequencing

DNA was extracted using the FastDNA SPIN® kit for Feces (MP Biomedical, Santa Ana, CA, U.S.; product: 116570200) by strictly following the user’s manual. The V4–V5 region of the 16S rRNA gene was amplified by PCR using the universal bacterial primers: 515-FB (GTGYCAGCMGCCGCGGTAA) and 926-R (CCGYCAATTYMTTTRAGTTT) (17, 39) following the cycling parameters in Thakkar et al. (29). This amplified product was cleaned using the AxyPrep Mag PCR Cleanup Kit (Corning, Inc., Corning, NY, U.S.), barcoded using the TruSeq dual indexing approach, and subsequently cleaned again, as previously described in detail (17). Barcoded amplicons were quantified using a Qubit dsDNA HS Assay Kit (Invitrogen, Carlsbad, CA, U.S.), and amplicons pooled. The Purdue Genomics Core Facility sequenced the pooled amplicons on an Ilumina MiSeq run with 2 × 250 cycles and V2 chemistry (Ilumina, Inc., San Diego, CA, U.S.), yielding 5,415,175 total reads.

### 2.4 Sequence analysis

16S rRNA gene amplicons were analyzed following the mothur MiSeq SOP (Kozich JJ Mothur Pipeline AEM 2013, accessed March 2018), using the following adaptations. The screen.seqs() command was used with a maximum homopolymer of 9 and a maximum length of 411 bp. Sequences were aligned against the mothur-formatted SILVA database (v.132) restricted to the V4-V5 region, and reads without alignment across positions 1970 – 15692 were removed. After removal of uninformative columns using filter.seqs(), sequences were pre-clustered using diffs=3 and checked for chimeras using VSEARCH (40) as implemented in mothur. Sequences were classified using the RDP Classifier (41, 42) as implemented in mothur using the RDP training set (to which species epithets had been added) as a reference (v. 16) at cutoff=95. Per-sample sequences were rarefied to 6,764 reads for α− and β−diversity analyses, resulting in a Good’s coverage above 97% for all samples. Diversity calculators implemented in mothur were used for α-diversity (species observed (sobs), Chao, Simpson evenness, Shannon, and inverse Simpson) and β-diversity (Bray-Curtis, Yue and Clayton’s theta, and Jaccard dissimilarity) analyses (43–48).

### 2.5 Metagenomics

#### 2.5.1 : Sequencing, Assembly, and Alignment

35 samples collected from various timepoints, sampling locations, and treatment size groups were sent to Purdue University’s sequencing core for 150bp paired-end shotgun sequencing using the Illumina NovaSeq platform (samples are listed in Supplemental Table 1) and analyses were executed on Purdue University’s Bell high performance computer cluster. Each sample (fastq) was checked for adapter content, contamination, and quality using bbmerge (v.39.00) (49) and fastQC (v0.11.99) (50). Two co-assemblies using SPAdes (v3.11.1 with -- meta flag) (51) were produced from the 35 samples being split into two groups: particle-associated plus controls and supernatant plus controls.

Each assembly was checked with MetaQuast.v3.2 (52) and reported an N_50_ of 9220 and 8948 for particle- and supernatant-associated assemblies, respectively. Both assemblies were independently indexed via Bowtie2 (v 2.3.5.1) (53) and each paired-end fastq file set was mapped to the index, compressed, and then sorted to create ordered BAM alignment files via Bowtie2. Anvi’o (v. 7.1) (54) was used to generate a contigs database for each assembly; BAM files mapping reads to each respective assembly were read in as a corresponding PROFILE.db. Read count and percent coverage for each sample-assembly combination was determined by HTSeq-count (v0.13.5) (55).

#### 2.5.2 : Binning and MAG Selection

Metagenomic binning was performed on both assemblies separately using MaxBin (v 2.2.3) (56), MetaBat (v 2:2.15) (57), and CONCOCT (v 1.1.0) (58). Read count data from HTSeq-count (v0.13.5) for each sample used in co-assembly (21 samples each) was used as inputs for each binning algorithm. DAS_Tool (v 1.1.2) (59) was used to analyze the outputs from the independent binning methods and combine them into one homogenous bin set. Bins from DAS_Tool with < 75% completion or > 25% redundancy were removed from further analysis. In total, the particle-associated assembly yielded 137 bins and the supernatant-associated assembly yielded 161 bins, hereafter referred to as metagenome-assembled genomes (MAGs). To reduce MAG redundancy across both assemblies, MAGs were merged if they passed an average nucleotide identity (ANI) threshold. fastANI (v1.32) (60) was run in a pairwise all-versus-all manner for all MAGs from both assemblies, resulting in a tab-delimited file comparing average nucleotide identity (ANI) and alignment fraction (aligning fragments of a query MAG to a reference MAG / total fragments of a MAG) of each MAG against every other MAG. Using an ANI threshold of 97% and an alignment fraction threshold of 0.95, we paired highly similar MAGs across metagenomic assemblies. We refer to these highly similar MAG pairs as analogs. Abundance profiles were analyzed for analogous MAGs, and pairs within 2% of each other for every sample were confirmed. 48 MAGs passed both ANI and abundance similarity thresholds and could be deemed true analogs of one another. The smaller (total nucleotide length) of the analogous pair was selected as the representative MAG resulting in a set of 252 MAGs across both assemblies.

Each MAG passed the gold-standard for MAG quality (2022 CAMI thresholds) and ranged in size from ∼1.25 million bases (mb) to over 6mb in total nucleotide content, with a mean MAG size of 2.6mb (standard deviation of 0.79mb). Gene content ranged from just over 1,000 to over 6,000 genes, with a mean of 2,443 genes (standard deviation of 694 genes).

#### 2.5.3 : Annotation Pipeline and Naming Convention

A custom snakemake (v 5.10.0) (61) file was created to reformat contigs, predict CDS regions, and annotate MAGs. Briefly, the snakemake script: reformatted all MAG fasta files to adhere to Anvi’o (v7.1) (54) internal rules, predicted MAG taxonomy using CATBAT (v5.2.3; metagenomic mode and default parameters) (62), predicted CDS regions using Prodigal (v2.6.3) (63), and annotated predicted CDS regions using hidden Markov models (HMMs) with HMMER (v3.3.2) (64) and the TIGRFAM (65), KOALA (66), Pfam (67), RAST (68), and COG (69) databases. Contig abundance and coverage results from 2.5.1 were used to calculate normalized per-MAG mean abundance (analogous to reads per kilobase per million base pairs (RPKM) as previously employed to detect MAG abundance changes) and coverage statistics (70). To normalize, first the total number of reads in a sample (specific fastq pair) was divided by 1 million to get a per-million normalization scalar. Then, total raw reads mapped to each contig was divided by the per-million scalar and subsequently divided contig length (in kilobases).

Each of the 252 MAGs was named by concatenating the two highest levels of annotation determined by CATBAT (separated by a “_”) and appending the assembly of origin (p or s, separated by a “-“). If the assembly resulted from a merged analogous pair, an asterisk was appended after the “p” or “s” designation. Finally, all MAGs were alphabetically sorted and a prefix from 1-252 was added (separated by a “_”).

#### 2.5.4 : MAG Association

MAGs were assigned to categories based on the average RPKM of contigs as calculated above. MAG RPKM values were first averaged across sample categories, (e.g., for sampling location into one of three categories: supernatant, particle, and initial). These average RPKM abundances were then compared across categories. If a MAG had an abundance 1.5 times greater in one category compared to others, it was assigned to the high abundance category. If the MAG RPKM values were within a 1.5x variance across all categories, it was not assigned to any.

Upon assigning MAGs, the KO and CAZY copy numbers annotated as above were analyzed with LEfSe (v.2017-02-03) to identify genes relevant for specific functions (71). KOs with an LDA (log 10) score > 2 across all sample groups were collapsed and a subset of these KOs extracted for further analysis. The percent of MAGs belonging to family *Lachnospiraceae* containing KOs related to flagella and pili formation was determined in Excel. CAZyme substrates was determined through the CAZy database (http://www.cazy.org, access date: 07/20/2023). Heatmap of KO presence across MAGs and LEfSe output was drawn in R (v4.1.1).

## 3. RESULTS

### 3.1 Wheat bran sugar composition is particle size-dependent

Here, unlike in our previous study (17), reduced particle sizes were generated from the same (the coarsest) parent brans, however chemical differences among fractions may still be introduced by biases in milling and sieving processes. To determine how particle size influenced WB fiber structures, neutral monosaccharide compositions were determined (Supplemental Figure 1A). We observed a correlation between WB particle size and the percent xylose present (R^2^=0.86), where the smallest fraction had less (38.96%) and the largest fraction had more (48.24%) across all fractions. In contrast, arabinose displayed an inverse correlation (R^2^=0.61), where the smallest size had the highest (41.42%) and the largest size had the lowest (34.21%) abundance. Although the ratio of arabinose to xylose generally decreased with increasing size, one of the intermediate size fractions (850-1000 um) had a high arabinose: xylose ratio, and a plateau was observed at the large size fractions (1000 -1700 um and >1700 um) (Supplemental Figure 1B). These data suggested that the arabinoxylans in the coarse bran fraction were less branched with arabinofuranosyl substituents than those in fine bran fractions.

### 3.2 Short-chain fatty acid production is dependent on wheat bran size

After fecal fermentation for 6, 12, 24, 36, and 48 hours, SCFAs, gas, and pH were determined as primary metabolic outputs. Fermented WB fractions did not perceptibly differ in either gas production or pH reduction (Figure 1). Although there were no differences in acetate concentrations or total SCFAs, propionate and butyrate production linearly related to WB fraction size; propionate was higher and butyrate lower in fine WB particle fermentations as compared to coarse. These particle size effects began to develop as early as six hours after inoculation, plateaued after 24 hours, and were most pronounced after 48 hours of fermentation. These differences in SCFAs implied altered microbial community metabolism, which may have been driven by the differences in component polysaccharide structures (e.g., arabinose: xylose ratio) or by differences in microbial access to and usage of the available carbohydrates in the physical matrix.

**Figure 1:**
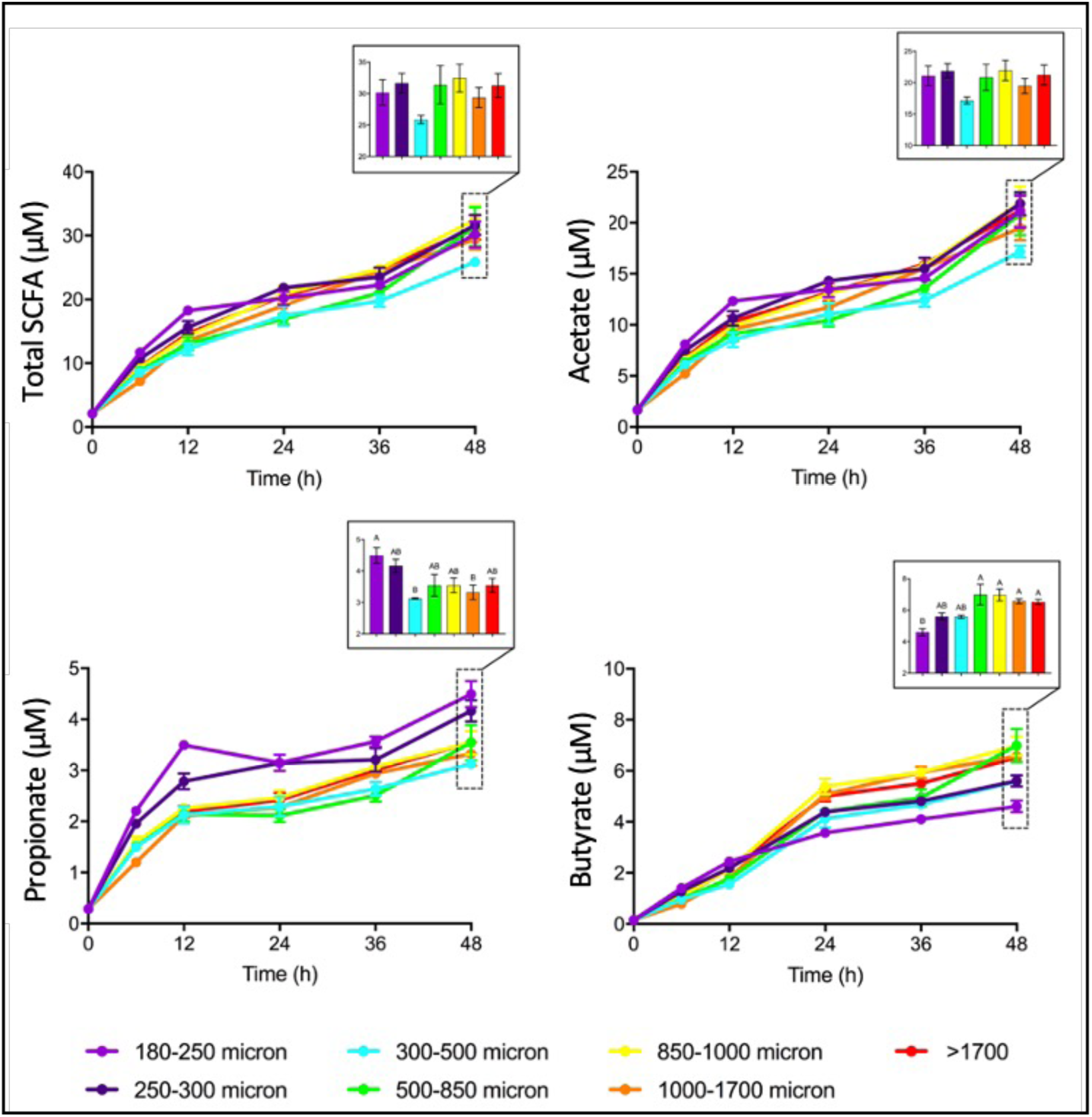
Fermentation of small wheat bran particles results in higher propionate and lower butyrate production compared to large wheat bran particles. Short-chain fatty acid (SCFA) production by fecal microbiota in *in vitro* fermentations over time with insets representing profiles at 48 hours. Total SCFA is the sum of acetate, propionate, and butyrate. Mean values with the same letter are not significantly different as determined by Tukey’s multiple comparisons test, p < 0.05. Error bars represent the standard error of the mean of three separate replicates.

### 3.3 Large and small wheat bran particles select for distinct taxa

To elucidate differences in the microbial composition after fermentation, microbes were collected separately from both the supernatant and the WB particles at 48 hours and the 16S rRNA genes were sequenced. Sequences that shared 97% identity were condensed into operational taxonomic units (OTUs; computational analogs of species). We calculated α-diversity (inclusive of richness, evenness, and combined metrics) for all WB fraction fermentations separately for supernatant and particle-associated organisms (Supplemental Figure 2). Although no discernable differences were observed for supernatant-associated fractions, higher richness + evenness combined metrics noted for smaller particle-attached communities than those attached to larger ones. Notably, the positive fermentation control, FOS, exhibited significantly lower α-diversity, indicating strong growth of a relatively small number of organisms on this substrate.

Communities colonizing fine and coarse bran particles exhibited distinct successional trajectories. Community β-diversity was also calculated to link bran particle size or association with community composition. At 12 hours, the supernatant-(circle) and particle-(square) associated communities were obviously distinguished but, within these categories, particle size was not determinative (Figure 2). At 24 hours, the small particle size shifted marginally, where the large particle size shifted drastically. Finally, at 48 hours, the samples re-clustered within the first two principal coordinates. This pattern was observed in both the supernatant- and particle-associated samples.

**Figure 2:**
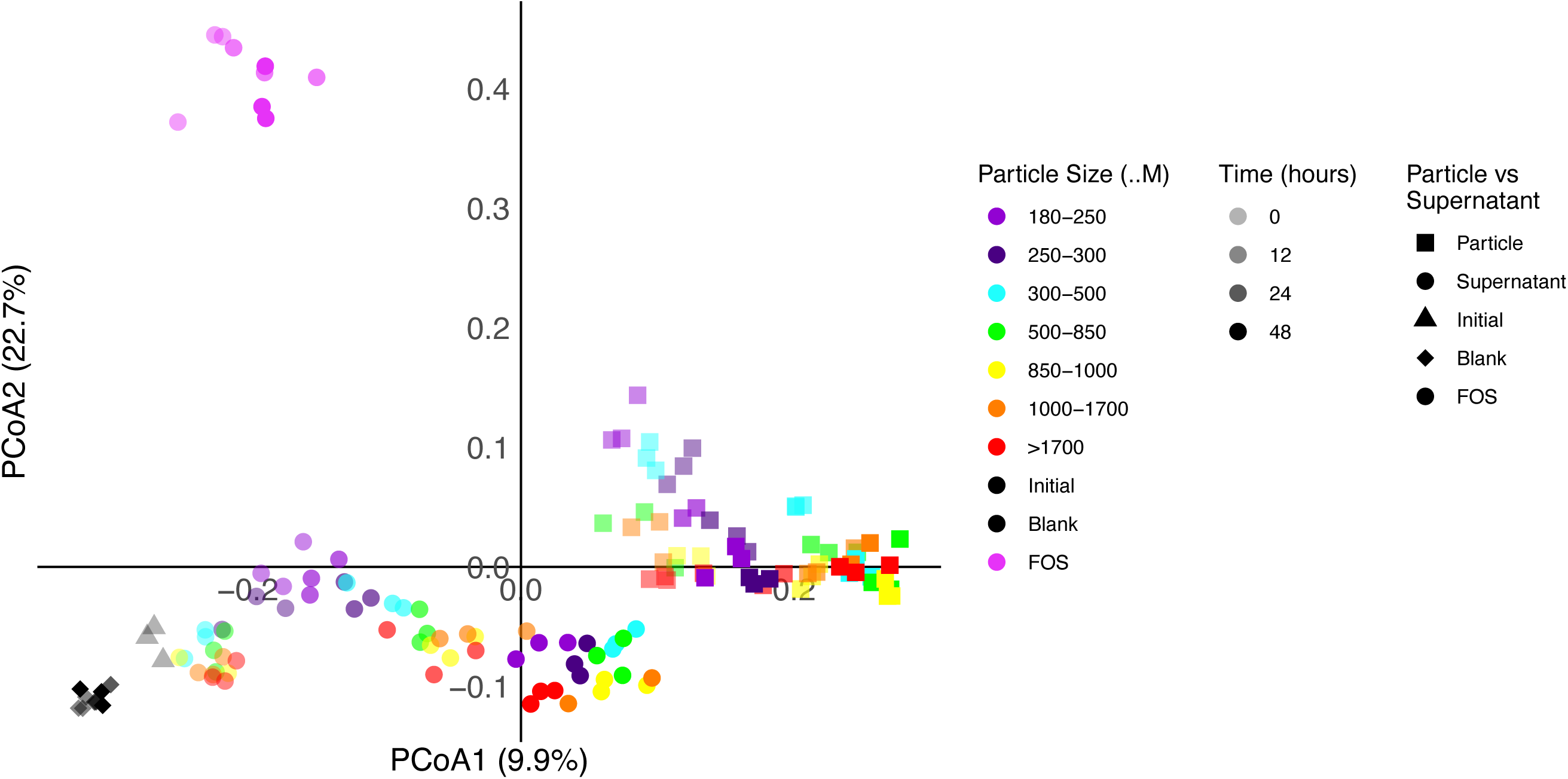
Community structures of particles and supernatant converge from 24 to 48h. Principal coordinate analysis of community structures associated with wheat bran size fractions, as determined by Bray-Curtis metric of 16S rRNA gene amplicon sequencing. FOS (fructooligosaccharide) was used as a fast-fermenting positive control, indicated by pink circles. Blank samples did not contain any substrate, indicated by black diamonds. Supernatant (circle), particle (square), 12h (high opacity), 24h (medium opacity), 48h (low opacity), colors are ordered by size, where violet = 180-250, indigo = 250-300, blue = 300-500, green = 500-850, yellow = 850-1000, orange = 1000-1700, red ≥ 1700.

The organisms responsible for driving divergent size-dependent shifts in β-diversity were classified within two main families, *Bacteroidaceae* and *Lachnospiraceae*, which contained individual members whose relative abundances shifted greatly (Figure 3). Some OTUs classified within the same genus exhibited drastically different abundances across the sample types. These abundance patterns suggest functional differences between closely related organisms that allow them to occupy the distinct niches afforded by variously sized bran particles.

**Figure 3:**
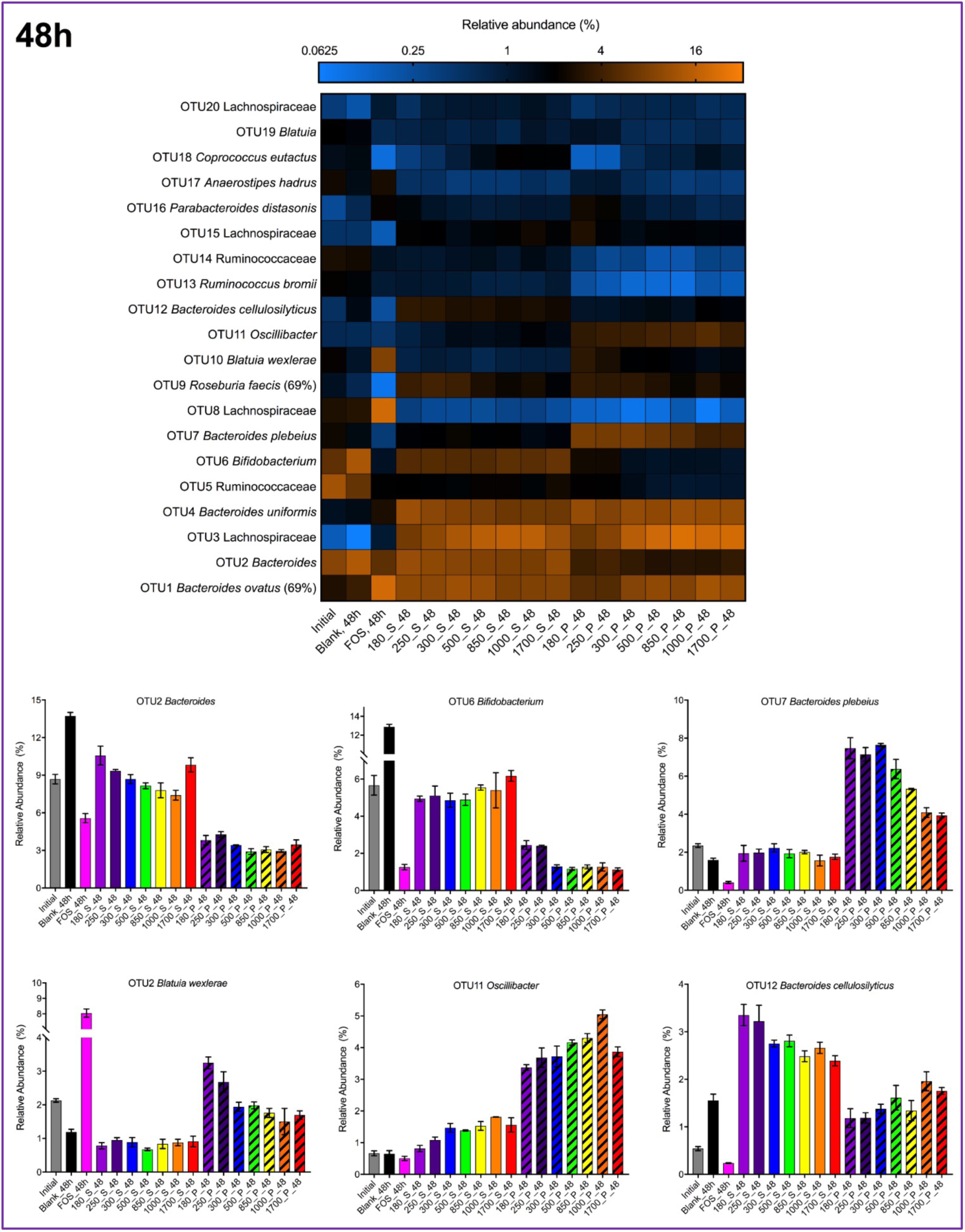
Individual taxa display relative abundance patterns associated with specific features. Relative abundances (percentage of sequences) of the top 20 OTUs in each sample are shown after fermentation for 48 hours.

### 3.4 Metagenome assembled genomes mostly belong to Bacillota

A heavily curated list of 252 MAGs was used for all downstream analyses (Supplemental Table 2, Supplemental Figure 3). The majority of MAGs were annotated as belonging to Bacillota (formerly, Firmicutes) (79%) with the remaining MAGs split between Bacteroidota (8%), Actinobacteriota (6%), Pseudomonadota (3%), Desulfobacterota (1%), and Verrucomicrobiota, Fusobacteriota, and Methanobacteria (≤ 1%) (Table 1). Families *Lachnospiraceae*, *Oscillospiraceae*, and *Ruminococcaceae* composed 31%, 15% and 14% of Bacillota MAGs, respectively.

**Table 1:** MAG associations. Most MAGs belonging to the same taxa cluster together in the associations with a location or time but not size. For Family, other represents taxa with 5 or fewer MAGs. In each sampling category, the relative abundance of each MAG was determined and categorized if the abundance was 1.5 times greater in each other category listed and (Location and Size) compared to the initial fecal sample. MAGs that failed to meet these criteria are placed in the “other” category.

| Family | Total | Location |  |  | Time |  |  |  | Size |  |  |  |  |  |
| --- | --- | --- | --- | --- | --- | --- | --- | --- | --- | --- | --- | --- | --- | --- |
|  | MAGs | Particle | Supernatant | Other | 0h | 24h | 48h | Other | Blank | FOS | Small | Medium | Large | Other |
| Acutalibacteraceae | 19 | 0 | 3 | 16 | 14 | 0 | 2 | 3 | 0 | 0 | 2 | 0 | 2 | 15 |
| Anaerovoracaceae | 12 | 0 | 6 | 6 | 4 | 0 | 2 | 6 | 0 | 2 | 5 | 3 | 0 | 2 |
| CAG-74 | 11 | 0 | 0 | 11 | 5 | 0 | 3 | 3 | 1 | 0 | 0 | 0 | 0 | 10 |
| Eggerthellaceae | 11 | 0 | 5 | 6 | 6 | 0 | 0 | 5 | 7 | 0 | 0 | 0 | 0 | 4 |
| Lachnospiraceae | 62 | 8 | 1 | 53 | 34 | 6 | 2 | 20 | 3 | 13 | 7 | 11 | 8 | 20 |
| Oscillospiraceae | 29 | 2 | 8 | 19 | 7 | 0 | 6 | 16 | 2 | 2 | 4 | 4 | 0 | 17 |
| Rikenellaceae | 7 | 0 | 0 | 7 | 0 | 0 | 0 | 7 | 2 | 0 | 0 | 0 | 0 | 5 |
| Ruminococcaceae | 28 | 4 | 4 | 20 | 10 | 1 | 5 | 12 | 2 | 1 | 7 | 4 | 1 | 13 |
| other | 73 | 4 | 17 | 52 | 19 | 1 | 18 | 35 | 21 | 3 | 19 | 13 | 1 | 16 |

After assembly and binning, MAGs were annotated with TYGS to identify the closest isolate, resulting in 122 MAGs for which there was a high-similarity near-neighbor isolate (dDDH_d4 > 70%) annotation and 130 MAGs for which there was a poor (dDDH_d4 < 70%) or missing annotation.

Of the well-described MAGs, the representation across phyla was similar to overall patterns (Bacillota 81%; Bacteroidota 8%; Actinobacteriota 5%; Pseudomonadota 4%; Desulfobacterota, Verrucomicrobiota, Fusobacteriota, and Methanobacteria at less than 1%). Of the remaining 130 MAGs for which there was no high similarity near-neighbor isolate, the taxonomic distribution was similar (Bacillota 77%; Bacteroidota 9%; Actinobacteriota 8%; Pseudomonadota 2%; Desulfobacterota 2% ; Verrucomicrobiota 2% ; and Fusobacteriota, and Methanobacteria at less than 1%); these potentially represent “dark matter” of the human gut microbiota.

### 3.5 Genes related to motility are associated with bran particle size

To ascertain which genomic features were associated with particle attachment and particle size, MAG relative abundances (as RPKM) were used to determine overrepresentation in each sample type. Initial comparisons identified only 16 MAGs associated with supernatant or particles at 24h or 48h, suggesting significant growth on WB, whereas the majority were associated with time zero (49 MAGs), or were neither associated with particles nor supernatant (97 MAGs). The 16 MAGs represented multiple members of *Lachnospiraceae* and *Oscillospiraceae*, with single representatives from other families (Supplemental Table 2).

An analysis of patterns across families indicated that some families were associated with supernatant fractions (*Acutalibacteraceae*, *Anaerovoracaceae*, *Eggerthellaceae*, and *Oscillospiraceae*), others were associated with particle fractions (*Lachnospiraceae*), and others had equal number of MAGs associated with both conditions (*Ruminococcaceae*). Similarly, some families demonstrated a preference for fractions containing small and medium sizes (*Anaerovoracaceae*, *Oscillospiraceae*, *Ruminococcaceae*) whereas others appeared to be equally distributed (*Lachnospiraceae*). We also noted that single MAGs representative of diverse families (*Acutalibacteraceae, Anaerovoracaceae*, *CAG-74*, and *Oscillospiraceae*) increased on bran particles between 0h and 48h, suggesting that the ability to colonize and grow on bran particles may be resolved at the species or strain level. The relative abundance of multiple MAGs belonging to *Lachnospiracea*e and *Oscillospiraceae* shifted greatly over time (Figure 4). An unbiased approach using LEfSe to identify genes significantly associated with sample types (localization, size, or sampling) across all taxa uncovered that a set of KEGG Orthology (KO) groups associated with motility were more prevalent in MAGs associated with large particles (Figure 5); in addition, these MAGs were enriched in genes involved in sugar utilization, amino acid metabolism, and cell regulation. However, KOs associated with localization or time (Supplemental Figure 4). Similarly, we found no differences in KO composition between MAGs with or without near neighbors. Closer inspection of specific genes related to attachment, flagella, and pilus formation in MAGs within one well-distributed family, *Lachnospiraceae*, indicated medium and large fractions had a greater number of MAGs containing genes related to biosynthesis of flagella, but not pili (Supplemental Table 3).

**Figure 4:**
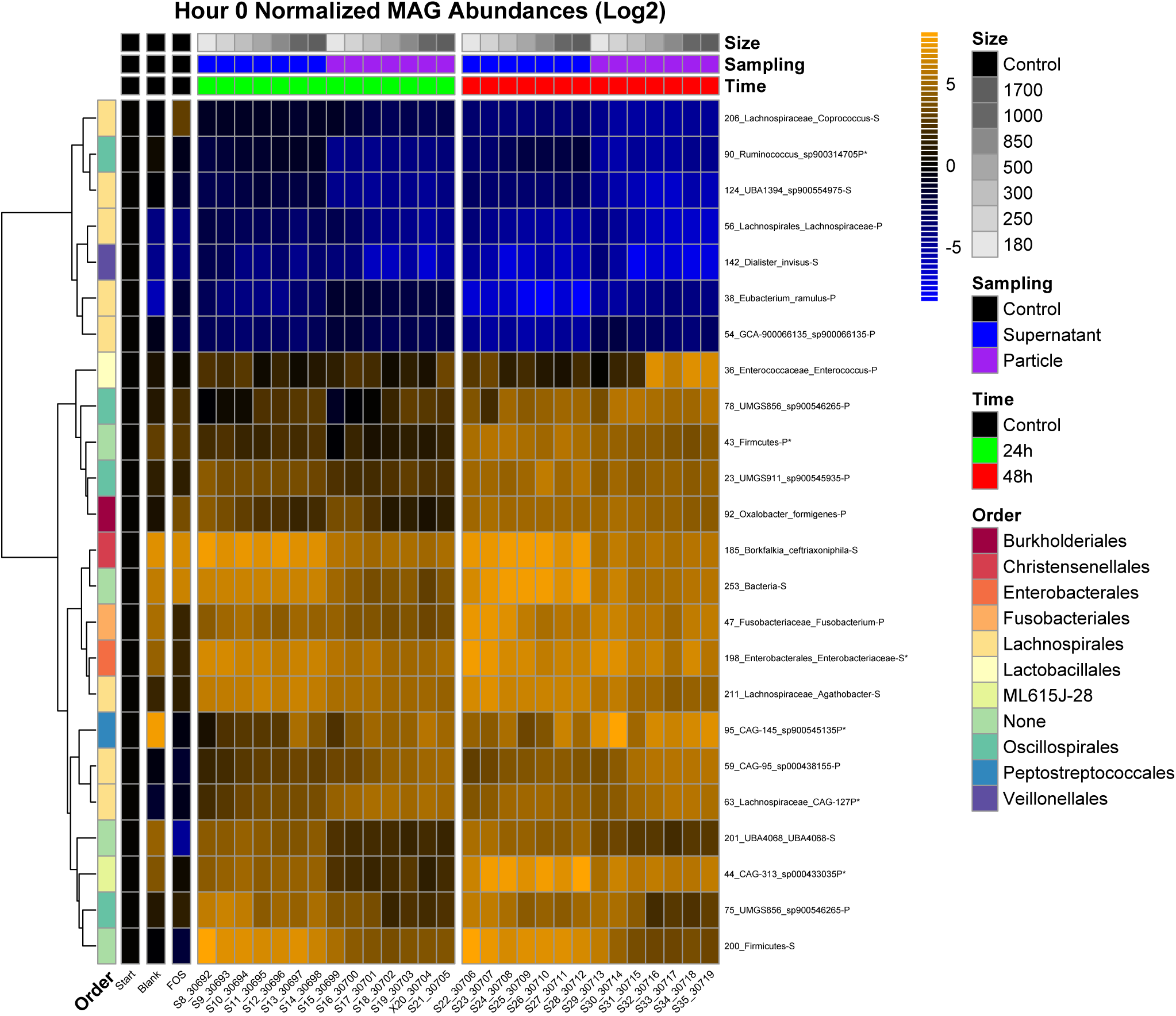
Individual MAGs display relative abundance patterns associated with specific features. Normalized relative abundances of the top 24 MAGs with the greatest change from the initial community are shown. Clustering by Euclidean distance indicates samples group primarily by sampling and time.

**Figure 5:**
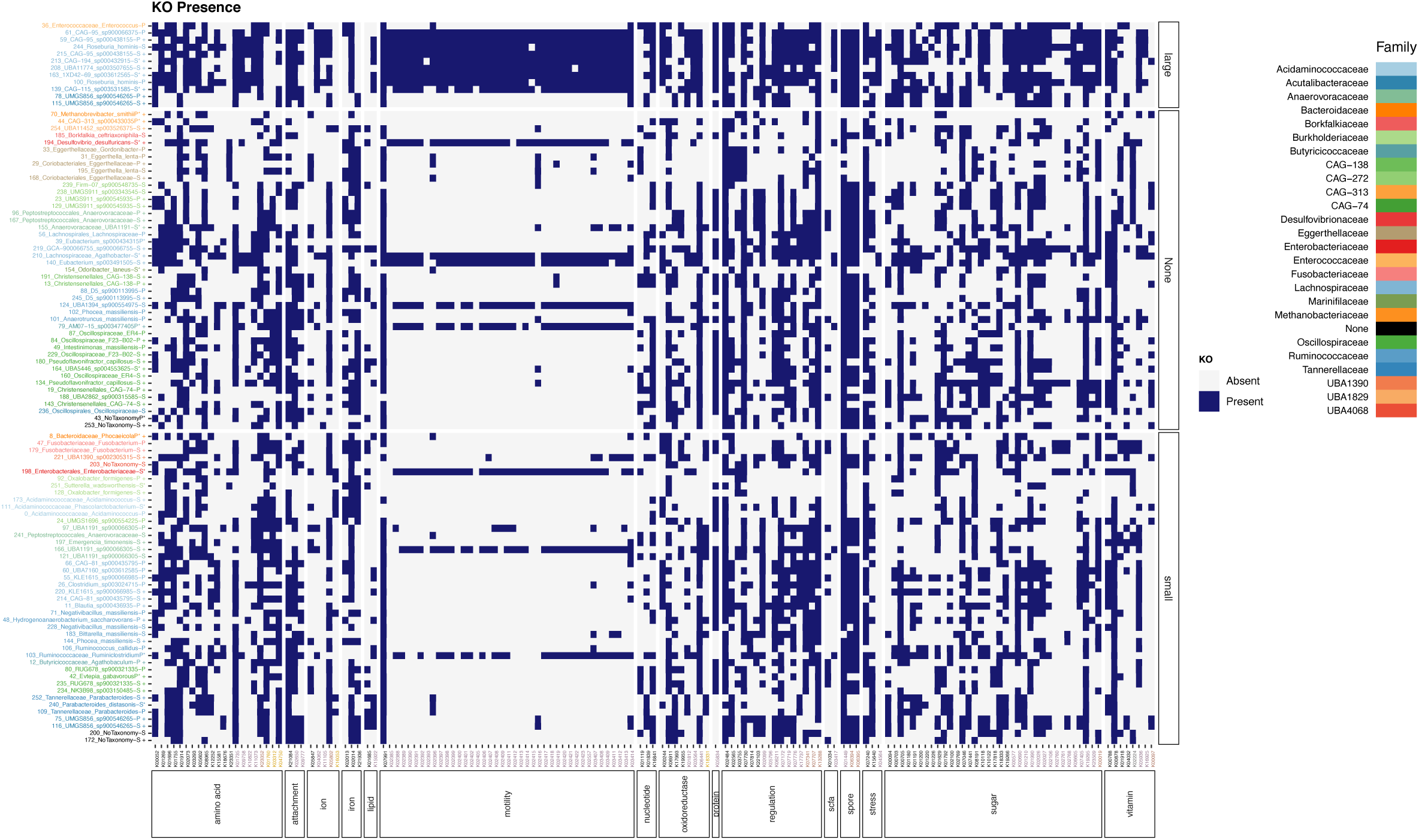
Organisms that contain genes related to motility and attachment are more abundant in large particles. Presence of genes of interest that discriminate for particle size, sampling time, particle association, or taxonomic family are faceted by pathway association. Organisms which had a poor (dDDH_d4 < 70%) or missing annotation through TGYS are indicated by a plus sign. Taxonomy is colored by family. Normalized relative abundance of mapped reads is used to facet MAGs into associated groupings.

### 3.6 Particle-associated MAGs are related to amylose and arabinoxylan degradation

KOs present in a majority of the MAGs in each category did not reveal any patterns or pathways that would indicate obvious differences in the metabolic niches afforded by the different particle sizes (Supplemental Table 4). However, a comparison of CAZymes revealed striking differences in the repertoire of these enzymes across MAGs associated with distinct localization, particle size, or incubation time. Myriad genes encoding putative amylases, amylomaltoses, N-acetylglucosaminidases, and lysozymes were overrepresented in particle-associated MAGs (Supplemental Table 5). No obvious enrichment patterns were observed for CAZymes present in supernatant-associated MAGs.

We then compared the CAZyme content of all MAGs using LEfSe (Figure 6, Supplemental Figure 5). Here, we also generally observed a greater number of glycoside hydrolases associated with MAGs enriched on particles (n=28) as compared to MAGs enriched in supernatant fractions (n=4) (Figure 6). The CAZymes associated with particle-associated MAGs were dominated by predicted amylases (GH126, GH13_20, GH13_31, GH13_11, GT35, and GH31) and genes targeted to linkages abundant in arabinoxylans (GH43_31, GH120, GH10, GH43_12, GH43_4, GH43_35, GH43_26, CBM22, GH127, GH36), supporting the role of these enzymes in WB fiber hydrolysis.

**Figure 6:**
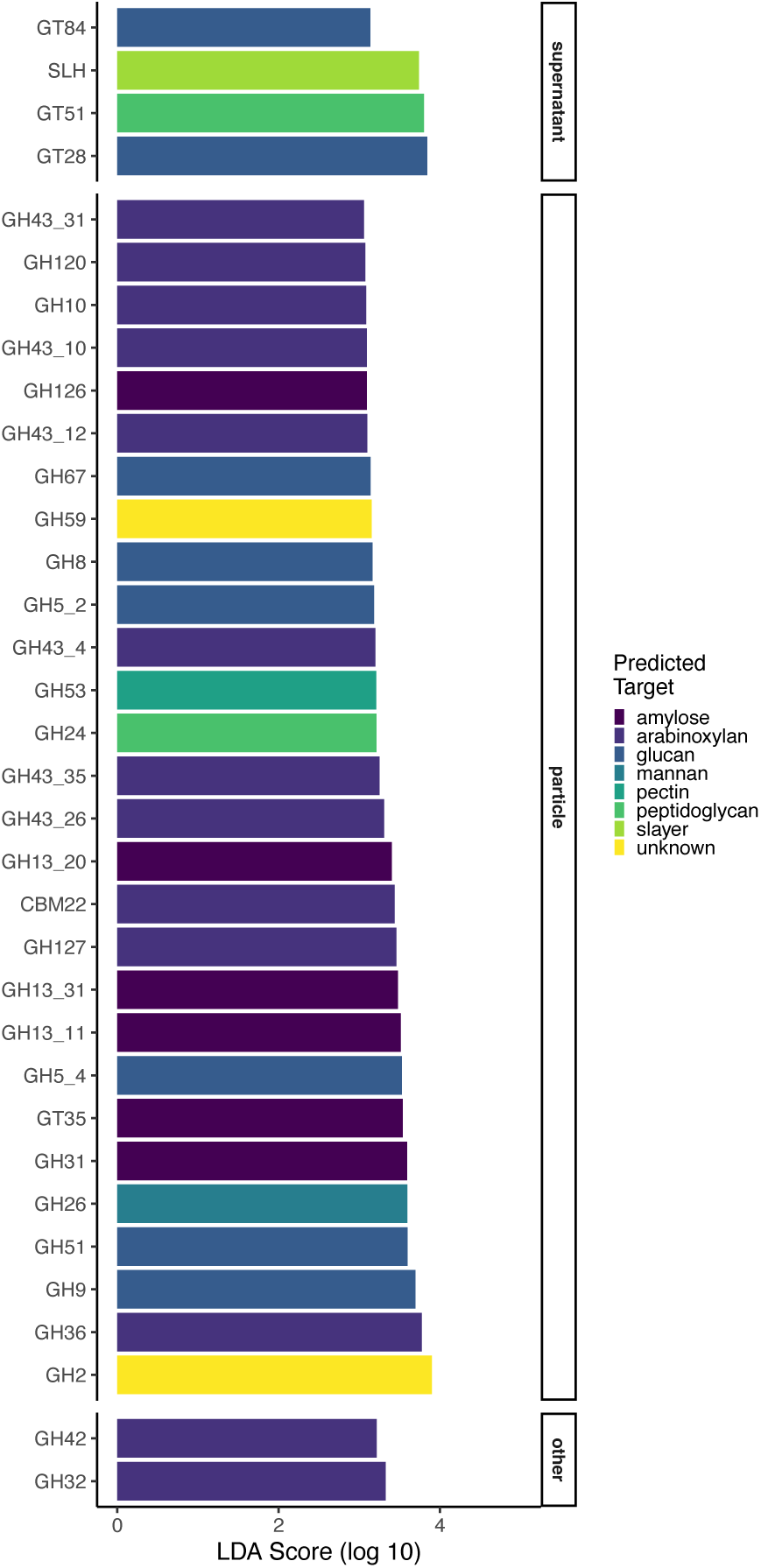
Differences in glycoside hydrolase abundances. A greater number of CAZymes are associated with particles. Within the particle-associated CAZymes, two enzymes families are dominant. Text color indicates glycoside hydrolase family.

A similar comparison across sizes revealed greater number of CAZymes in large WB fractions (n=10) as compared small (n=0) or medium (n=2) WB fermentations (Supplemental Figure 5). The CAZymes from MAGs significantly associated with large WB fractions included multiple predicted arabinoxylan-directed CAZymes (GH43_26, GH43_5, CE2, and GH51).

## 4. DISCUSSION

Cereal-derived dietary fibers are predominantly consumed in complex mixtures in the form of grain flours, in which the heterogeneous deposition of fibers within the plant cell (largely, its cell wall) limits the bioavailability of any specific polysaccharide and, therefore, requires liberation by successions of microbial enzymes prior to degradation and fermentation (31, 32, 72). Research on the impact of finely-resolved dietary fiber structures on gut community composition points to differences in organisms’ abilities to compete and cooperate in their degradation, internalization, and metabolism (16, 19). However, the large differences in spatial patterning of dietary fibers across a bran particle may strongly influence colonization efficiency across individual taxa (17, 72). As such, WB particles may be best regarded ecologically as patchy resources (73). Here, we used batch fecal fermentation on a range of sizes of bran particles generated by cyclone milling to identify bacterial taxa and genomic signatures driving differences in colonization and growth patterns on divergent WB fine structures.

We first demonstrated that WB particles generated from a single batch vary in their neutral sugar content, suggesting different polysaccharide structures and potentially distinct niches. Supporting this hypothesis, metabolic output during growth revealed a trade-off between propionate and butyrate production that trended with size. Concurrently, we also noted that community structures were particle-size dependent; specifically, that different OTUs within the same genus exhibited distinct attachment and growth patterns in size-dependent ways. Metagenomics revealed genes and associated metabolic functions that may be responsible for these growth differences. Notably, motility and capacity to directly metabolize particle polymers (i.e. putative amylases and xylanases) were overrepresented in large particle-associated MAGs at 24h. These data point to two important points: 1) genotypic variation of microbial taxa influences their access to different polysaccharides (74–77) and 2) structural context and spatial patterning of fibers in the diet (e.g., enmeshed with an insoluble matrix or in solution) likely affects the rate at which they are liberated and accessed by host microbiota (72, 78). Differences in rates of polysaccharide degradation, in turn, alter primary degrader fermentation and shape the pool of substrates available to secondary consumers, both with respect to liberated carbohydrates and metabolic products (e.g., lactate) that may be further metabolized by other microbes (79–82). Assignment of microbes to putative primary and secondary consumption modes will require measurement of gene expression patterns; however, we submit that lack of particle attachment suggests increased probability that microbiota operate largely in a secondary consumption mode. Given that differently sized particles vary in underlying polysaccharide structure and that OTU and MAG abundances vary with particle attachment and size preferences, different species (especially, *Bacteroides* spp. often observed in WB fermentations) diverge in their direct access to bran polysaccharides in the matrix and, potentially, in the fine polysaccharide structures they target (83).

Differences between the particle-associated and supernatant-associated microbial communities in this batch system contrast with similarities between these communities reported previously for continuous flow fermenter systems (84). Divergent succession observed before 24h between particle and supernatant communities is likely due to simple (mono- and di-) saccharide release by particle-associated microorganisms, as has been observed for complex polysaccharides (85, 86). Smaller WB fractions also likely possess a larger colonizable area and therefore result in increased growth of cross-fed organisms. For example, *Bifidobacterium* species have previously been reported to grow in the supernatant surrounding WB particles through crossfeeding (35) despite some species of *Bifidobacterium* lacking secreted glycoside hydrolases (76). Reduction of acetate and lactate produced by bifidobacteria may, in turn, support the growth of secondary consumers by serving as electron sinks, contributing to high overall SCFA production (17, 87). Similarly to our previous study on size-dependent WB fermentation, we also noted lower levels of butyrate in small WB fractions compared to medium and large WB particles (17). In our previous study, however, we sieved particles directly from mill fractions; here we create the size range of particles from identical (large) parent brans. Thus, differences in particle chemical composition are likely derived from preferential breakage patterns that give rise to compositional differences (88, 89). Others have observed strong inter-individual differences in WB fermentation (35, 72, 90) that drive individual shifts in metabolic output on different size WB fractions (91). Here, the microbial inoculum was derived from multiple donors to avoid deficiency of specific microbes within any one individual’s gut microbiome and provide for consistency in particle colonization. However, this artificially high diversity may have driven similarities between late-stage supernatant and particle communities, potentially increasing the rate of nutrient liberation and the exhaustion and dissociation of microbiota from the particle (92, 93), allowing slower-fermenting organisms or cross-fed organisms to ‘catch up’ (94).

Our metagenomic approach also reveals genomic characteristics of microbiota colonizing variously sized WB particles, advancing our understanding of WB fermentation through reconstruction of genomes of under-characterized organisms. Many unique MAGs that were associated with particles or supernatants at later time points (24h and 48h) are of interest for their potential in degradation of WB and have few or no sequenced phylogenetic neighbors. Multiple MAGs we recovered had the same taxonomic classifications but unique KO and CAZyme profiles. Although these may represent distinct strains of the same organism (95), since the genomes are not closed, the differences between these MAGs does not definitively indicate their distinct classification or gene content (96). However, when considering all overrepresented genomes associated with sample categories, we identified an abundance of putative starch metabolism-related genes, (e.g., amylases and amylomaltases), in particle-associated MAGs. These data may suggest the presence of residual starches following *in vitro* upper GI digestion, as has been described previously (36). We also observed a greater number of genes, such as N-acetylglucosidases and lysozymes, that might be implicated in attachment by altering surface charge or biofilm formation on particles, allowing for enhanced attachment or out-competition of early-attaching organisms (97). Whereas others have noted increased colonization of WB particles by members of the *Lachnospiraceae* (90), our study suggests that colonization differences among *Lachnospiraceae* are likely due to species-specific differences in functional capacity (95). Specifically, we noted genes related to motility in this family associated with larger particle sizes. Similar effects are observed in other ecosystems; genes associated with flagella and chemotaxis have been associated with marine particles (98). Our observations, however, might be greatly impacted by the incubation conditions in which higher shaking velocity may select for the growth of highly motile organisms on fast-sinking particles (99) and absence of exogenous nutrients may select against auxotrophs (18). Further, batch cultivation cannot disentangle differences in spatial organization that may impact colonization dynamics (100). It Future studies should examine size-dependent microbiota growth patterns under continuous-flow systems that more closely model *in vivo* dynamics (101).

Bioavailability of dietary nutrients is often constrained due to entrapment within particles, requiring liberation by microbial enzymes (31, 32, 72). Initial particle colonizers can resist colonization by subsequent microorganisms (33, 102, 103). The forces that drive colonization may include social interactions, such as attachment; nutrient acquisition, such as polysaccharide metabolism; metabolic interactions; quorum sensing; and growth (104). Understanding the genes relevant to colonization may reveal dietary control points for rational manipulation of fibers to shape the gut microbiome towards human health. In this study, we demonstrate that differently sized bran fractions generated from an identical source have different resource availability and ferment to different metabolic and community composition outcomes. We also find distinct ecological roles for different organisms despite similar or identical genome content, pointing to species- or strain-level ecotypes adapted for distinct WB niches. Further, we find that genes related to motility are more tightly associated with WB colonization than genes related to polysaccharide fermentation, suggesting that motility phenotypes may dictate initial colonization and fermentation of particulate substrates in the large intestine. Overall, these findings point to the importance of milling – both method and the particle sizes produced – which results in differences in fine structures and resource availability on the growth and metabolism of gut microbial species.

## 5. ACKNOWLEDGEMENTS

S.R.L designed and conceived the overall study. A.D.F., D.G.D., and C.W.E. analyzed the metagenomic data. Y.E.T. and R.T. analyzed the chemical and amplicon data. Y.E.T, R.T, and A.R.M. conducted the fecal fermentations. S.R.L supervised the research and received funding. A.D.F., D.G.D., and S.R.L. wrote the first draft of the manuscript. All authors approved the final manuscript.

---

**Supplemental Figure 1:** Sugar composition of wheat bran particles is unique between small and large particle sizes. Composition of neutral monosaccharides including rhamnose, arabinose, xylose, mannose, and glucose (A) and the arabinose to xylose ratio (B) of wheat brans fractions. Mean values with the same letter are not significantly different as determined by Tukey’s multiple comparisons test, p < 0.05. Error bars represent the standard error of the mean of three separate replicates.

**Supplemental Figure 2:** Community diversity and richness is higher in supernatant fractions and on small particles. Community membership across a range of particle sizes for the supernatant and particle fractions after 48 hours of fermentation. Jitter plot of all three replicates for each sample group for the number of OTUs (A) and Shannon Index (B) are shown.

**Supplemental Figure 3:** Genome size and total gene count for each MAG (A). Completion and contamination for each MAG (B). Number of MAGs per Phylum (C) and Order (D); MAGs without Order- or Phylum-level annotations are omitted.

**Supplemental Figure 4:** Organisms that contain genes related to motility and attachment are more abundant in large fractions. Presence of genes of interest that discriminate for sampling time (A) or particle association (B) are faceted by pathway association along the X axis. Organisms which had a poor (dDDH_d4 < 70%) or missing annotation through TGYS are indicated by a plus sign. Taxonomy is colored by family. Normalized relative abundance of mapped reads is used to facet MAGs into associated groupings.

**Supplemental Figure 5:** Differences in glycoside hydrolase abundances. A greater number of CAZymes are associated with 24h, and large sizes. Within the particle-associated CAZymes, two enzymes families are dominant. Text color indicates glycoside hydrolase family.

**Supplemental Table 1:** Sample information including code, wheat bran particle size, size category, sampling time, and sample type.

**Supplemental Table 2:** MAG statistics. Information includes per-sample relative abundance, TYGS analysis and taxonomy, categorization, total length, number of contigs, N50, and GC_content. MAG names are also included.

**Supplemental Table 3:** Copy number of genes involved in flagella and pili assembly in the Lachnospiraceae family across samples. Table includes percent of each gene across related samples.

**Supplemental Table 4:** Kegg Ontology (KO) counts for each MAG across samples.

**Supplemental Table 5:** Carbohydrate Active Enzymes (CAZy) counts for each MAG across samples.

